# Processivity of molecular motors under vectorial loads

**DOI:** 10.1101/2020.04.13.039784

**Authors:** Hamid Khataee, Zoltan Neufeld, Mohammed Mahamdeh

## Abstract

Molecular motors are cellular machines that drive the spatial organisation of the cells by transporting cargoes along intracellular filaments. Although the mechanical properties of single molecular motors are relatively well characterised, it remains elusive how the three-dimensional geometry of a load imposed on a motor affects its processivity, i.e., the average distance that a motor moves per interaction with a filament. Here, we theoretically explore this question for a single kinesin molecular motor by analysing the load-dependence of the stepping and detachment processes. We find that the processivity of kinesin increases with lowering the load angle between kinesin and microtubule filament, due to the deceleration of the detachment rate. When the load angle is large, the processivity is predicted to enhance with accelerating the stepping rate, through an optimal distribution of the load over the kinetic transition rates underlying a mechanical step of the motor. These results provide new insights into understanding of the design of potential synthetic biomolecular machines that can travel long distances with high velocities.

## 1 Introduction

Kinesin, dynein, and myosin molecular motors exert localised forces on intracellular components by stepping directionally along intracellular filaments using the energy released from the hydrolysis of adenosine triphosphate (ATP) [1]. The motors have fundamental roles in several cellular processes [1]. For instance, they mediate the changes in and maintenance of the morphology of cells, by directed transport of certain organelles [2] and sliding filaments along each other [3]. They also contribute to the movement of the cell body, by transporting the migration signals and controlling the cell polarity [4–6]. Accordingly, defects in the function of molecular motors cause various diseases, such as neurodegeneration and cancer [7, 8]. Therefore, a comprehensive understanding of the mechanical properties of molecular motors would provide new insights into the roles the motors play in health and disease.

A major factor that influences the function of a molecular motor in a three-dimensional (3D) cellular environment is the load applied on the motor, due to, for example, blockages caused by other cellular components or thermal fluctuations of the cargo carried by a motor [9–12]. The 3D vectorial character of the load ***F*** = (*F*_*x*_, *F*_*z*_) was established in experiments by Howard and colleagues [13]. They measured the vertical load component *F*_*z*_, where the horizontal component *F*_*x*_ was resisting (< 0, applied against the stepping direction); see Fig. 1. More recently, single-molecule optical trapping experiments [14–17] have studied the function of molecular motors under assisting loads *F*_*x*_ (> 0, applied in the stepping direction) as well; see Fig. 1. They have shown that the motors exhibit similar responses to the applied load direction: i) their velocity decreases with resisting loads, but changes slightly when the load is assisting, and ii) motors favor faster detachment rate under assisting loads than resisting ones. The importance of this directionality comes from the need to understand the collective function of a complex of multiple motors under a 3D load, which usually occurs within cells [18]. In such a complex, individual motors experience either resisting or assisting load due to intermolecular interactions.

**Figure 1:**
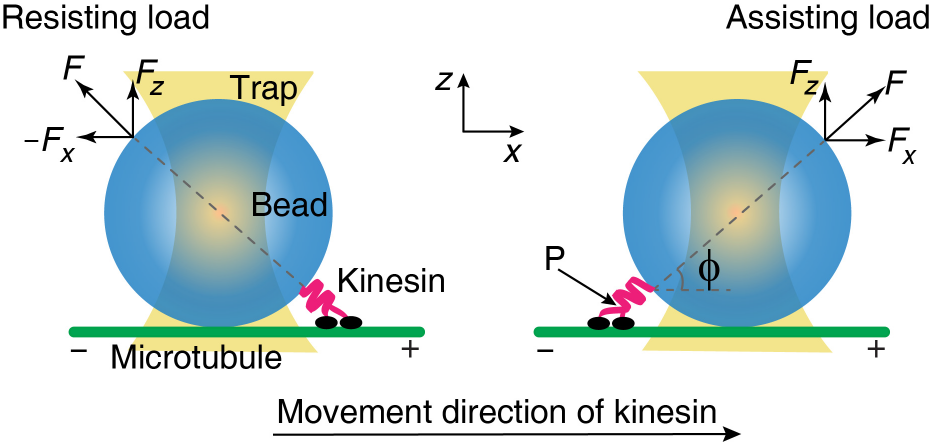
A graphical illustration of a microtubule-kinesin-bead complex in a typical optical trap assay (approximately to scale). Kinesin is attached to the bead (blue) with its tail (magenta) and moves, with its heads (black), along the microtubule (MT, green) toward the plus-end under a resisting (left) or an assisting (right) load transmitted to the point P on the motor, which links its tail to its heads, with an angle *ϕ*. Horizontal and vertical load components *F*_*x*_ and *F*_*z*_ are applied parallel and perpendicular to the MT axis, respectively.

The 3D vectorial character of load was originally quantified by Fisher and Kolomeisky [19, 20]. They modelled single-kinesin load-velocity data [13, 21] by quantifying the load-dependence of the mechanical kinetics underlying a stepping process of the motor [19, 20]. Recently, we also modelled the mechanics of force generation by kinesin motors by quantifying the detachment process of the motor as a function of an applied load ***F*** vector [22]. These models of the stepping and detachment processes as functions of the geometry of an applied load suggest kinesin motor as a model system for studying the 3D load-dependence of the processivity of molecular motors (i.e., the average distance that a motor moves per interaction with a filament [23–25]), which remains elusive.

Here we model the processivity of a single kinesin as a function of a 3D applied load vector. We determine the processivity using kinetic transition rate models that illustrate the displacement of the motor body in the (*x, z*) domain during stepping and detachment processes. The model describes how the vectorial load-dependencies of stepping and detachment processes determine the processivity.

## 2 Results and discussion

The processivity of a single kinesin motor with a mean run length

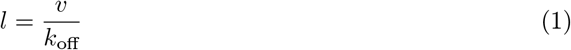

is maximised if the velocity *v* of the motor is high and the detachment rate *k*_off_ of the motor from a microtubule (MT) is low. This means that the relationship between the processivity and the vectorial character of the applied load depends on the role of the load in two processes: i) stepping, which determines the velocity *v*, and ii) detachment, which determines the run time (1/*k*_off_, average time to detachment). Therefore, to model the load-processivity relationship, we analyse the load-velocity and load-detachment rate relationships. We use Block and coworkers experimental data [14] as it provides measurements of velocity, detachment rate, and run length under resisting and assisting loads.

First, the effect of the vectorial character of the load on the velocity is described using a two-state transition rate model developed for an individual stepping cycle of the motor [19, 26]. This transition rate model describes a forward step of length *d* = 8.2 nm as passing through a sequence of two mechanochemical states 0 (ATP-free) and 1 (ATP-processing) [19]:

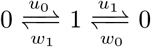

One complete forward transition corresponds to a forward step of the motor and the hydrolysis of an ATP molecule to its products, adenosine diphosphate (ADP) and inorganic phosphate (P_i_). Likewise, one complete reverse transition includes a backward step and the synthesis of ATP from ADP and P_i_ [20, 27]. The forward and backward transition rates are given by [19, 20]:

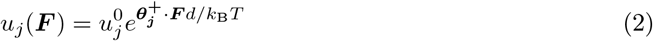

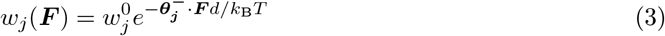

where 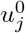 and 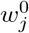 are the unloaded rates, 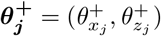 and 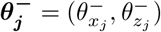 are the dimensionless load sharing factors (for *j* = 0, 1), and *k*_B_*T* is the thermal energy. The velocity is then calculated as [20]:

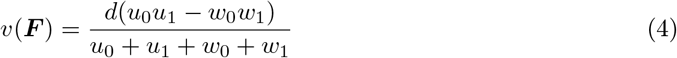

The parameter values of Equations (4) have been estimated by fitting to the experimental data of Block and colleagues [21]. The unloaded rates take 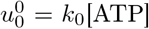, where *k*_0_ = 1.35 μM^−1^s^−1^, 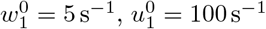, and 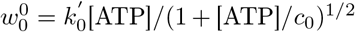, where 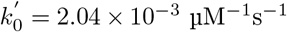 and *c*_0_ = 20 μM [19]. The force distribution vectors in the (*x, z*) plane are as [19]:

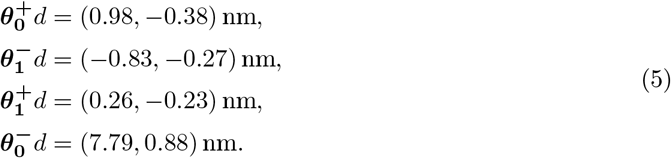

where the load distribution factors 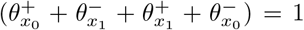 and 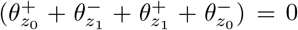 demonstrate the motion of the attachment point P in the (*x, z*) domain; see Fig. 2(A). They also indicate how the load is shared between the forward and backward rates according to the location of the kinetic transition states (i.e., the saddle points in the free-energy landscape) between states 0 and 1 [19, 27]. The transition states before and after ATP binding are located at 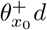 and 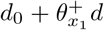 along the MT axis, respectively [19]. Using this two-state stepping transition rate model, it was found that the velocity generally decreases with increasing *F*_*z*_ [19, 26], rather than increases as argued in the MT buckling experiments [13]. It was reasoned that in the MT buckling experiments kinesin moves away from a curved (stressed) region of MT and that curvature of the MT might affect the motility of kinesin [26].

**Figure 2:**
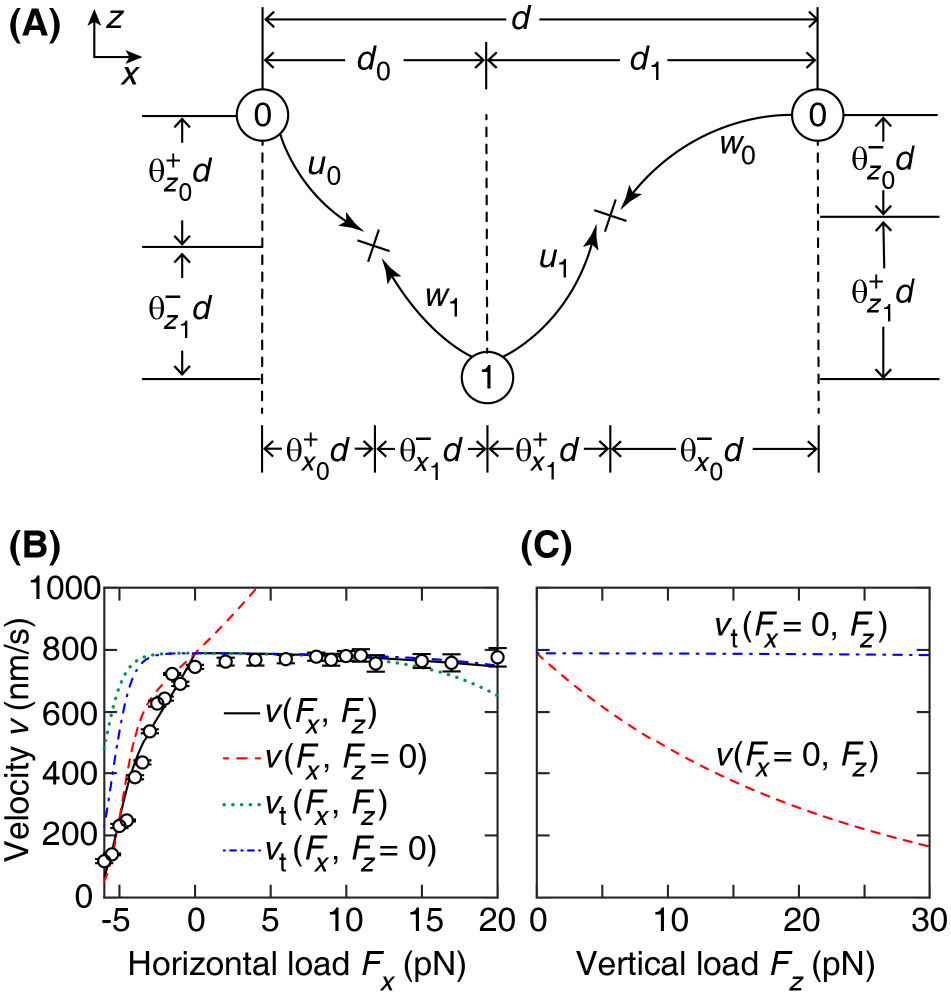
Velocity of single kinesin. (A) Schematic of the two-state stepping transition rate model from [19]. States 0 and 1: ATP-free and ATP-processing states, respectively. Crosses: transitions states. (B) Velocity *v* versus the horizontal load component *F*_*x*_. Open circles: experimental data (mean SE) with a 440-nm-diameter bead at saturating 2 mM ATP [14]. Solid curve: *v* where the motor is subject to both *F*_*x*_ and *F*_*z*_, calculated using Equation (4). Dashed curve: *v* when the load ***F*** is horizontal. Dotted curve: *v*_*t*_ denotes the velocity *v* when 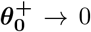 and 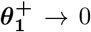. Dashed-dotted curve: *v*_*t*_ when the force ***F*** is horizontal. (C) *v* versus the vertical load component *F*_*z*_ (dashed curve). Dashed-dotted curve: *v*_*t*_.

The location of a transition state was also found to affect the velocity: kinesin moves faster when a transition state is close to the initial state [27–30]. The applied load increases the height of the kinetic barriers, making it more difficult to transition between states, and thus slowing down the velocity of the motor [28]. Therefore, if the distance to a transition state is zero, the load does not increase the height of the kinetic barrier along a mechanical step of the motor.

We analyse the effects of both the load angle *ϕ* and the locations of the transition states on velocity. We model the load-dependent velocity data [14] using Equation (4) and the ansatz *F*_*z*_ = *F*_*x*_tan*ϕ*; see Fig. 2(B), solid curve. We find that, for horizontal loads, the velocity increases, especially when the load is assisting; see Fig. 2(B), dashed curve. But, for vertical loads, velocity decreases; Fig. 2(C), dashed curve. We then find that the velocity increases under all load geometries when the distance to the transition states approaches zero, i.e., 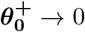 and 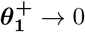; see Fig. 2(B), dotted and dashed-dotted curves and Fig. 2(C) dashed-dotted curve. Together, these results suggest that velocity increases under various applied load geometries with lowering the distance to transition states. In this regime, an enhancement in velocity is predicted under resisting loads. For vertical loads, velocity remains almost load-independent.

To model the effect of the vectorial character of the load on the detachment process, we follow our model [22] that describes the detachment of a kinesin from a MT as a two-step process, which passes through three states:

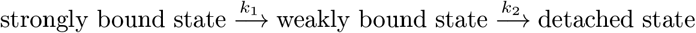

*k*_1_ and *k*_2_ are the fast and slow detachment rates, respectively, that depend on the load ***F*** and displacement 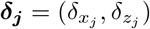 vectors according to 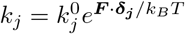, where 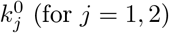 are the unloaded rates. The effective detachment rate is then given by [22]:

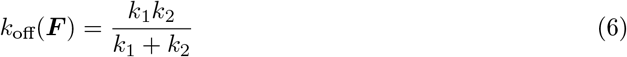

By fitting Equation (6) to the load-detachment rate data [14], the two-step detachment model described the data by a continuous curve, where 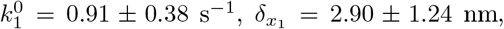 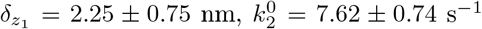, and 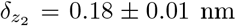(mean ± SE) [22]. This two-step model explained the effects of the load geometry on the detachment process [22]: i) detachment rate decreases with horizontal loads (i.e., catch-bond behaviour) when the force is load, and increases for assisting loads (see Fig. 3(A), dashed curve); and ii) detachment rate increases with vertical loads (i.e., slip-bond behaviour); see Fig. 3(B). Using these catch and slip bonding mechanisms, we explained different behaviours of kinesin motors reported from different laboratories [22]. It explained that the stall force of multiple kinesin can be similar to that of a single kinesin when the vertical load component is high, as observed in [31, 32], due to slip-bond behaviour. However, multiple kinesins can produce forces much larger than the single-motor force when the vertical load component is small, e.g., in gliding assays [33, 34] and bead assays [35–37], due to catch-bond behaviour. This later prediction has been recently confirmed in a three-bead assay where lower detachment rates were observed for a single kinesin with low kinesin-MT angle [38].

**Figure 3:**
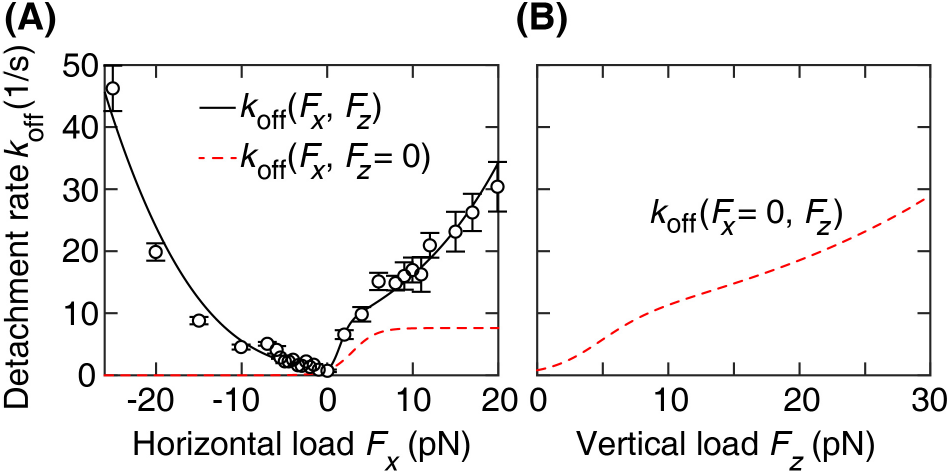
Detachment rate of single kinesin. (A) Detachment rate *k*_off_ versus *F*_*x*_. Open circles: experimental data (mean ± SE) from [14]. Solid curve: *k*_off_ calculated using Equation (6), where the motor is subject to both *F*_*x*_ and *F*_*z*_. Dashed curve: *k*_off_ when the applied load ***F*** is parallel to the MT axis. (B) *k*_off_ when ***F*** is perpendicular to the MT axis.

We now model the processivity of kinesin by considering the load-dependencies of the stepping and detachment processes. We find that the processivity increases over all resisting and assisting loads with lowering the kinesin-MT angle; see Fig. 4(A). This is explained by the catch-bond behaviour of the motor and the acceleration of the motor velocity; see Figs. 2(B) and 3(A). In addition, decreasing the distance to the stepping kinetic transition states, enhances the run length over resisting loads; see Fig. 4(A). This is due to the increase in the velocity of the motor. For vertical loads, lowering the distance to the stepping kinetic transition states is predicted to benefit the run length; see Fig. 4(B). Small distances to transition states results in an almost steady velocity, while motor exhibits slip-bond behaviour; see Figs. 2(C) and 3(B).

**Figure 4:**
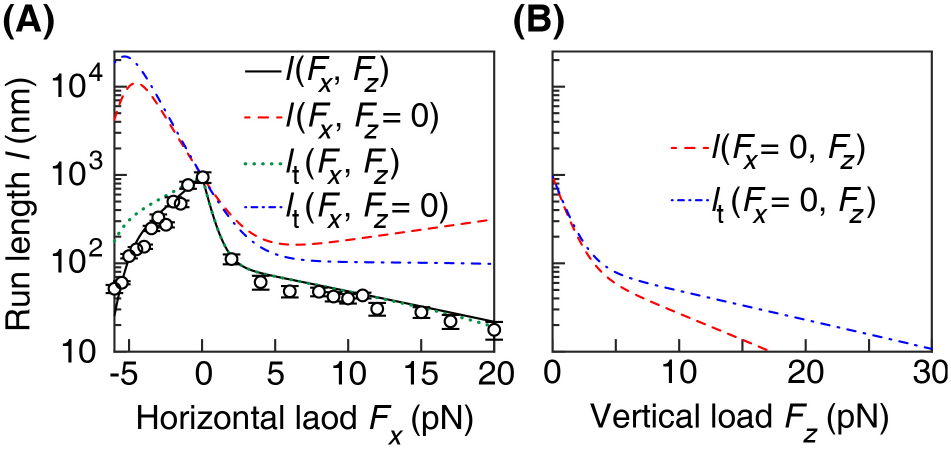
Run length of single kinesin. (A) Run length *l* versus *F*_*x*_. Open circles: experimental data (mean SE) from [14]. Solid curve: *l*, where the motor is subject to both *F*_*x*_ and *F*_*z*_, derived using Equation (1). Dashed curve: *l* when the force ***F*** is horizontal. Dotted curve: *l*_*t*_ denotes the run length when 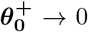 and 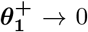. Dashed-dotted curve: *l*_*t*_ when the force ***F*** is horizontal. (B) *l* versus *F*_*z*_ (dashed curve). Dashed-dotted curve: *l*_*t*_.

We conclude that a motor with longer body length (for a fixed size of cargo), where distances to stepping transition states are small, can travel long distances with high velocities; see Figs. 2(B) and 4(A), dashed-dotted blue curves. Such motors exhibit catch-bond behaviour with faster stepping rates. On the other hand, run length of a motor is minimised when the motor body is short and distances to stepping transition states are high. For such motors, stepping rates are slow and slip-bond behaviour is more likely to occur. An *in vivo* implication of this prediction is that motors with different mechanical properties can exhibit different processivities in carrying cargoes with different sizes.

In this study, we modelled the processivity of a single kinesin motor under 3D applied load geometries by taking into account the load-dependent dynamics of the stepping and detachment processes. We found that the processivity of kinesin increases with lowering the vertical component of the load. This is explained by the catch-bond behavior that allows kinesin to stay attached to MT under high loads *F*_*x*_. This behaviour has been observed in recent experiments where the effects of horizontal and vertical loads on the attachment duration of a single kinesin were investigated using a three-bead assay [38]. When kinesin was subject to horizontal forces, threefold increase in the attachment duration was observed. Run length of kinesin under loads with different vertical components can be measured by running similar experiments in force clamp mode analogous to experiments performed on myosin motors [15, 39, 40]. Our model also predicted that, when the vertical load component is large, processivity is enhanced with accelerating the stepping rate, through minimising the load distribution over the stepping kinetic rates.

Future work can also consider the effects of the backward stepping (velocity) of the motor on its processivity, upon the availability of the velocity and detachment rate data over high resisting and assisting loads. Backward steps of kinesin, on average, are driven by ATP hydrolysis at force above the stalling conditions [10, 41–44], although ATP synthesis can result in backward stepping [19, 20]. It is interesting to study the effect of the ATP concentration on the processivity. Although Block and colleagues have measured the ATP dependence of the run length under moderate assisting force (4 pN) [45], this would require a comprehensive load- and ATP-dependent detachment rate data.

The present model can further be extended to study the effects of the vectorial character of the applied load on the mechanics of dynein and myosin motors, since the load-velocity and -detachment rate behaviours of kinesins, dyneins, and myosins are similar [14–17]. This can help to understand the mechanics of collective function of different molecular motors, owing to the force-dependent interactions of the motors. This will provide new insights into exploring the design of synthetic molecular machines powered by molecular motors [46] that can operate with high efficiency in a 3D environment.

## Acknowledgement

We are grateful to Jonathon Howard for valuable comments on an early version of the results.

## Notes

### Competing Interest Statement

The authors have declared no competing interest.

### Summary of Updates

We reviewed the text.

## References

1 J. Howard, Mechanics of motor proteins and the cytoskeleton (Sinauer, Sunderland, MA, USA, 2001).

2 V. I. Rodionov, “Microtubule-dependent control of cell shape and pseudopodial activity is inhibited by the antibody to kinesin motor domain”, J. Cell Biol. 123, 1811–1820 (1993).

3 A. L. Jolly, H. Kim, D. Srinivasan, M. Lakonishok, A. G. Larson, and V. I. Gelfand, “Kinesin-1 heavy chain mediates microtubule sliding to drive changes in cell shape”, Proc. Natl. Acad. Sci. USA 107, 12151–12156 (2010).

4 J.-B. Manneville, M. Jehanno, and S. Etienne-Manneville, “Dlg1 binds gkap to control dynein association with microtubules, centrosome positioning, and cell polarity”, J. Cell Biol. 191, 585–598 (2010).

5 A. Bachmann and A. Straube, “Kinesins in cell migration”, Biochem. Soc. T. 43, 79–83 (2015).

6 R. B. Vallee, G. E. Seale, and J.-W. Tsai, “Emerging roles for myosin II and cytoplasmic dynein in migrating neurons and growth cones”, Trends Cell Biol. 19, 347–355 (2009).

7 A. J. Lucanus and G. W. Yip, “Kinesin superfamily: roles in breast cancer, patient prognosis and therapeutics”, Oncogene 37, 833–838 (2018).

8 S. T. Brady and G. A. Morfini, “Regulation of motor proteins, axonal transport deficits and adult-onset neurodegenerative diseases”, Neurobiol. Di. 105, 273–282 (2017).

9 M. J. Schnitzer, K. Visscher, and S. M. Block, “Force production by single kinesin motors”, Nat. Cell Biol. 2, 718–723 (2000).

10 N. J. Carter and R. A. Cross, “Mechanics of the kinesin step”, Nature 435, 308–312 (2005).

11 A. Gennerich, A. P. Carter, S. L. Reck-Peterson, and R. D. Vale, “Force-induced bidirectional stepping of cytoplasmic dynein”, Cell 131, 952–965 (2007).

12 A. D. Mehta, R. S. Rock, M. Rief, J. A. Spudich, M. S. Mooseker, and R. E. Cheney, “Myosin-v is a processive actin-based motor”, Nature 400, 590–593 (1999).

13 F. Gittes, E. Meyhöfer, S. Baek, and J. Howard, “Directional loading of the kinesin motor molecule as it buckles a microtubule”, Biophys. J. 70, 418–429 (1996).

14 J. O. Andreasson, B. Milic, G.-Y. Chen, N. R. Guydosh, W. O. Hancock, and S. M. Block, “Examining kinesin processivity within a general gating framework”, eLife 4, e07403 (2015).

15 L. Gardini, S. M. Heissler, C. Arbore, Y. Yang, J. R. Sellers, F. S. Pavone, and M. Capitanio, “Dissecting myosin-5b mechanosensitivity and calcium regulation at the single molecule level”, Nat. Commun. 9, 2844 (2018).

16 S. Can, S. Lacey, M. Gur, A. P. Carter, and A. Yildiz, “Directionality of dynein is controlled by the angle and length of its stalk”, Nature 566, 407–410 (2019).

17 Y. Ezber, V. Belyy, S. Can, and A. Yildiz, “Dynein harnesses active fluctuations of microtubules for faster movement”, Nat. Phys. 16, 312–316 (2020).

18 E. L. Holzbaur and Y. E. Goldman, “Coordination of molecular motors: from in vitro assays to intracellular dynamics”, Curr. Opin. Cell Biol. 22, 4–13 (2010).

19 M. E. Fisher and Y. C. Kim, “Kinesin crouches to sprint but resists pushing”, Proc. Natl. Acad. Sci. USA 102, 16209–16214 (2005).

20 M. E. Fisher and A. B. Kolomeisky, “Simple mechanochemistry describes the dynamics of kinesin molecules”, Proc. Natl. Acad. Sci. USA 98, 7748–7753 (2001).

21 S. M. Block, C. L. Asbury, J. W. Shaevitz, and M. J. Lang, “Probing the kinesin reaction cycle with a 2d optical force clamp”, Proc. Natl. Acad. Sci. USA 100, 2351–2356 (2003).

22 H. Khataee and J. Howard, “Force generated by two kinesin motors depends on the load direction and intermolecular coupling”, Phys. Rev. Lett. 122, 188101 (2019).

23 J. Howard, A. J. Hudspeth, and R. D. Vale, “Movement of microtubules by single kinesin molecules”, Nature 342, 154–158 (1989).

24 R. D. Vale, T. Funatsu, D. W. Pierce, L. Romberg, Y. Harada, and T. Yanagida, “Direct observation of single kinesin molecules moving along microtubules”, Nature 380, 451–453 (1996).

25 S. M. Block, L. S. B. Goldstein, and B. J. Schnapp, “Bead movement by single kinesin molecules studied with optical tweezers”, Nature 348, 348–352 (1990).

26 Y. C. Kim and M. E. Fisher, “Vectorial loading of processive motor proteins: implementing a landscape picture”, J. Phys. Condens Mat 17, S3821–S3838 (2005).

27 J. A. Wagoner and K. A. Dill, “Molecular motors: power strokes outperform brownian ratchets”, J. Phys. Chem. B 120, 6327–6336 (2016).

28 J. Howard, “Motor proteins as nanomachines: the roles of thermal fluctuations in generating force and motion”, in Biological physics (Springer Basel, 2011), pp. 47–59.

29 J. A. Wagoner and K. A. Dill, “Mechanisms for achieving high speed and efficiency in biomolecular machines”, Proc. Natl. Acad. Sci. USA 116, 5902–5907 (2019).

30 T. Schmiedl and U. Seifert, “Efficiency of molecular motors at maximum power”, Europhys. Lett. 83, 30005 (2008).

31 K. Furuta, A. Furuta, Y. Y. Toyoshima, M. Amino, K. Oiwa, and H. Kojima, “Measuring collective transport by defined numbers of processive and nonprocessive kinesin motors”, Proc. Natl. Acad. Sci. USA 110, 501–506 (2013).

32 D. K. Jamison, J. W. Driver, A. R. Rogers, P. E. Constantinou, and M. R. Diehl, “Two kinesins transport cargo primarily via the action of one motor: implications for intracellular transport”, Biophys. J. 99, 2967–2977 (2010).

33 V. Bormuth, A. Jannasch, M. Ander, C. M. van Kats, A. van Blaaderen, J. Howard, and E. Schäffer, “Optical trapping of coated microspheres”, Opt. Express 16, 13831 (2008).

34 A. Hunt, F. Gittes, and J. Howard, “The force exerted by a single kinesin molecule against a viscous load”, Biophys. J. 67, 766–781 (1994).

35 G. T. Shubeita, S. L. Tran, J. Xu, M. Vershinin, S. Cermelli, S. L. Cotton, M. A. Welte, and S. P. Gross, “Consequences of motor copy number on the intracellular transport of kinesin-1-driven lipid droplets”, Cell 135, 1098–1107 (2008).

36 M. Vershinin, B. C. Carter, D. S. Razafsky, S. J. King, and S. P. Gross, “Multiple-motor based transport and its regulation by tau”, Proc. Natl. Acad. Sci. USA 104, 87–92 (2007).

37 M. Bovyn, S. Gross, and J. Allard, “Molecular motor organization and mobility on cargos can overcome a tradeoff between fast binding and run length”, bioRxiv, 1–16 (2019).

38 S. Pyrpassopoulos, H. Shuman, and E. M. Ostap, “Modulation of kinesin’s load-bearing capacity by force geometry and the microtubule track”, Biophys. J. 118, 243–253 (2020).

39 M. Capitanio, M. Canepari, M. Maffei, D. Beneventi, C. Monico, F. Vanzi, R. Bottinelli, and F. S. Pavone, “Ultrafast force-clamp spectroscopy of single molecules reveals load dependence of myosin working stroke”, Nat. Methods 9, 1013–1019 (2012).

40 R. S. Rock, S. E. Rice, A. L. Wells, T. J. Purcell, J. A. Spudich, and H. L. Sweeney, “Myosin vi is a processive motor with a large step size”, Proc. Natl. Acad. Sci. USA 98, 13655–13659 (2001).

41 M. Nishiyama, H. Higuchi, and T. Yanagida, “Chemomechanical coupling of the forward and backward steps of single kinesin molecules”, Nat. Cell Biol. 4, 790–797 (2002).

42 H. Khataee and A. W.-C. Liew, “A stochastic automaton model for simulating kinesin processivity”, Bioinformatics 31, 390–396 (2015).

43 S. Liepelt and R. Lipowsky, “Kinesin’s network of chemomechanical motor cycles”, Phys. Rev. Lett. 98, 258102 (2007).

44 H. Khataee, S. Naseri, Y. Zhong, and A. W.-C. Liew, “Unbinding of kinesin from microtubule in the strongly bound state enhances under assisting forces”, Mol. Inf. 37, 1700092 (2018).

45 B. Milic, J. O. L. Andreasson, W. O. Hancock, and S. M. Block, “Kinesin processivity is gated by phosphate release”, Proc. Natl. Acad. Sci. USA 111, 14136–14140 (2014).

46 H. Hess, “Toward devices powered by biomolecular motors”, Science 312, 860–861 (2006).

